# Aperture: Accurate detection of structural variations and viral integrations in circulating tumor DNA using an alignment-free algorithm

**DOI:** 10.1101/2020.12.04.409508

**Authors:** Hongchao Liu, Huihui Yin, Guangyu Li, Junling Li, Xiaoyue Wang

## Abstract

**Background:** The identification of structural variations (SV) and viral integrations in circulating tumor DNA (ctDNA) is a key step in precision oncology that may assist clinicians for treatment selection and monitoring. However, it is challenging to accurately detect low frequency SVs or SVs involving complex junctions in ctDNA sequencing data due to the short fragment size of ctDNA.

**Results:** Here, we describe Aperture, a new fast SV caller that applies a unique strategy of *k*-mer based searching, breakpoint detection using binary labels and candidates clustering to detect SVs and viral integrations in high sensitivity, especially when junctions span repetitive regions, followed by a barcode-based filter to ensure specificity. We evaluated the performance of Aperture in stimulated, reference and real datasets. Aperture demonstrates superior sensitivity and specificity in all tests, especially for low dilution test, compared with existing methods. In addition, Aperture is able to predict sites of viral integration and identify complex SVs involving novel insertions and repetitive sequences in real patient data.

**Conclusions:** Using a novel alignment-free algorithm, Aperture achieves sensitive, specific and fast detection of structural variations and viral integrations, which may enhance the diagnostic value of ctDNA in clinical application. The executable file and source code are freely available at https://github.com/liuhc8/Aperture.

## Background

Circulating tumor DNA (ctDNA) is tumor-derived DNA fragment found in the bloodstream. Mutations in ctDNA have been identified as excellent biomarkers for early detection and treatment monitoring of cancer [1]. Although ctDNA was first described 50 years ago, until recently base-resolution analysis of ctDNA can be performed with next generation sequencing technology [1]. Targeted deep sequencing has allowed detection of multiple types of cancer-specific mutations in ctDNAs, such as single nucleotide variants, small insertion and deletions, and structural variations [1]. As ctDNA is only a small fraction of total cell-free DNA (cfDNA), to analyze sequence alterations in high sensitivity, several molecular barcoding-based targeted sequencing approaches have been developed, such as TEC-seq [2]. Background error-correction methods tailored for barcoded sequencing has enabled detection of single-nucleotide variants with fractions as low as 0.02% in cfDNA [3]. However, due to the short fragment size and extremely low fraction of ctDNA, accurate detection of structural variations in ctDNAs, especially for translocations and insertion/deletions that are larger than 50 bp, are still challenging.

Most existing algorithms identify SVs from next-generation sequencing (NGS) data based on alignment, such as GRIDSS [4], Lumpy [5], SvABA [6], DELLY [7] and CREST [8]. They inferred SVs from the reads or read pairs that are either mapped to two distant loci (split reads and discordant read pairs) or only partly mapped (soft-clip reads). SViCT has optimized the alignment-based strategies for cfDNA dataset with ultra-high read depth and extremely low dilutions [9]. However, although repetitive sequences are often substrates for recombination that leads to structural rearrangement of genome [10], most the alignment-based SV callers have difficulty in detecting complex SVs involving repetitive sequences using short reads [11]. Present alignment tools, such as BWA, randomly selects one of the possible locations in the genome for repetitive sequences and output them as low-quality alignments, which prevents SV callers to cluster the evidence together to infer the breakpoint. Similarly, SViCT is not compatible for SVs involving viral integrations, which could provide valuable information for cancer development and diagnosis. For example, HBV DNA integration near *TERT* promoter can result in telomerase overexpression, which is required for the proliferation of liver cells in hepatocellular carcinoma development [12].

Here we introduce Aperture, an alignment-free SV caller for cfDNA sequencing that considers complex SVs involving repetitive sequence and viral integrations (Fig.1). Starting from raw sequencing data in FASTQ format, Aperture performs a *k*-mer based database searching involving three different libraries and implements SV breakpoint detection by a fast approximation of set intersection using binary labels. Aperture then gathers candidate reads with identical junctions or similar genomic positions to achieve fault-tolerant evidence clustering for SV detection. The final output from Aperture includes predicted SV, number of supporting molecules, mapping quality of both breakends and sequences of identical micro-homology at breakpoints.

**Figure 1.**
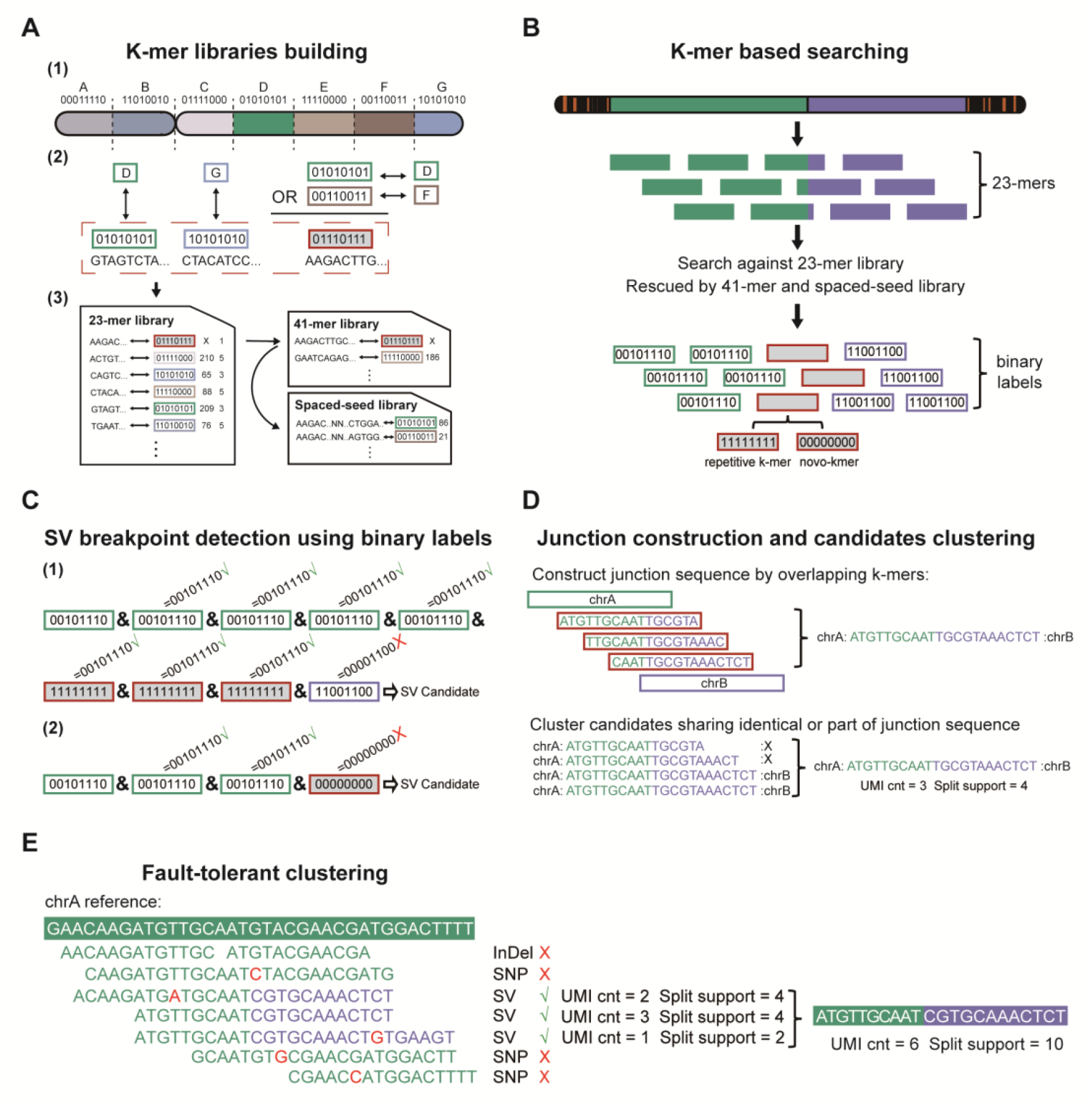
Overview of the Aperture structural variation detection tool for cfDNA dataset. (A) The workflow of building *k*-mer libraries: (1) Reference genome is divided into segments of at most 2,500 bp in length. Then, each segment is randomly assigned with a 32-bit binary label containing only five 1-bits. For clarity, binary labels in this figure are illustrated as a shorter version (8-bit) with only four 1-bits. (2) Binary labels of *k*-mers coming from multiple segments are recalculated using a bitwise OR operation. (3) 23-mers derived from reference genome are sorted and stored in 23-mer library along with their binary labels, offsets and scores. To enhance the precision of mapping, 41-mer library and spaced-seed library are constructed for genomic regions covering repetitive 23-mers. (B-E) The workflow of SV discovery: (B) 23-mers cut from cfDNA reads are searched against the 23-mer library to get their binary labels. If consecutive repetitive 23-mers are found in a read, 41-mers and spaced-seeds will be generated to perform remapping. There are two possible situations for *k*-mers spanning breakpoints. When the breakpoint covers repetitive region in genome, binary labels of *k*-mers spanning the junction are still valid but contain more than five 1-bits, otherwise they are not valid, representing a blunt junction. (C) A bitwise AND operation is performed iteratively on all binary labels in a read, and the validities of the resulting labels (i.e. contain at least five 1 bits) are checked to identify breakpoint candidates with homologous junctions (1) or blunt junctions (2). (D) Junction sequence is constructed by consecutive *k*-mers covering the breakpoints. Then, SV candidates sharing identical or part of junction sequence are clustered and merged. (E) In order to tolerate sequencing errors, PCR errors and mutations, candidates with similar genomic positions are clustered together. Also, the similarity of 3’ end of the breakpoint and reference is checked to avoid false positives result from variants near the breakpoint junction.

After performance test using simulated cfDNA data, we found that Aperture achieved much higher sensitivity and specificity than existing SV callers, at dilution ranging from 0.1% to 10%. We also applied Aperture on three real patient cfDNA datasets in different clinical settings. Apertured successfully detected druggable translocations in lung cancer patients and HBV integration in *TERT* promoter in liver cancer patients, including a complex rearrangement involving repetitive sequence that was missed by most SV callers. In addition, Aperture is efficient in CPU and runs extremely fast in parallel mode. Implemented in Java, Aperture is available as an open source tool for the reliable detection of SVs and viral integration in cfDNA dataset on all operating system, for both scientific research and clinical practice.

## Implementation

### Overview

Aperture is developed to achieve sensitive detection of breakpoints introduced by structural variations in cfDNA datasets. It is based on: (i) a unique strategy of *k*-mer based searching, which uses two different *k* lengths and spaced-seed to optimize the coverage of repetitive sequences at breakpoints (ii) a fast approximation of intersection approach to identify breakpoint junctions containing either novo-*k*mers or repetitive sequences and (iii) a barcode based filtering strategy designed for cfDNA dataset with molecular barcoding. The workflow of Aperture consists of the following steps: (1) Construction of a 23-mer library with repeat scores as well as binary labels representing genomic positions. The library is built once for each genome and serves as a reference library for searching against 23-mers derived from NGS reads. For those genomic regions that cannot be uniquely covered by 23-mers, a 41-mer library and a spaced-seed library are built (Fig.1A). (2) Due to the short fragment size of ctDNA (50-166 bp) [13], paired-end (PE) reads are merged together to give a single sequence. After PE reads merging, *k*-mers derived from NGS read are searched against three different libraries involving 23-mers, 41-mers and spaced-seeds[14] (Fig.1B). (3) After *k*-mer based searching, a fast set intersection of genomic regions, implemented by a bitwise AND operation of binary labels, is performed to identify potential SV breakpoints (Fig.1C). Breakpoint candidates with low qualities are filtered out. (4) Candidates with identical junctions or similar positions are clustered together, and sequence similarity of right break ends and reference is examined to avoid false positives result from sequencing errors, PCR errors and mutations (Fig.1D,E). (5) To further achieve high specificity, all candidates are evaluated according to *k*-mer mapping qualities as well as counts of supporting *k*-mers, NGS reads and barcodes. A summary of Aperture stages is shown in Figure 1 and details for each step are provided in the following sections.

**Figure 2.**
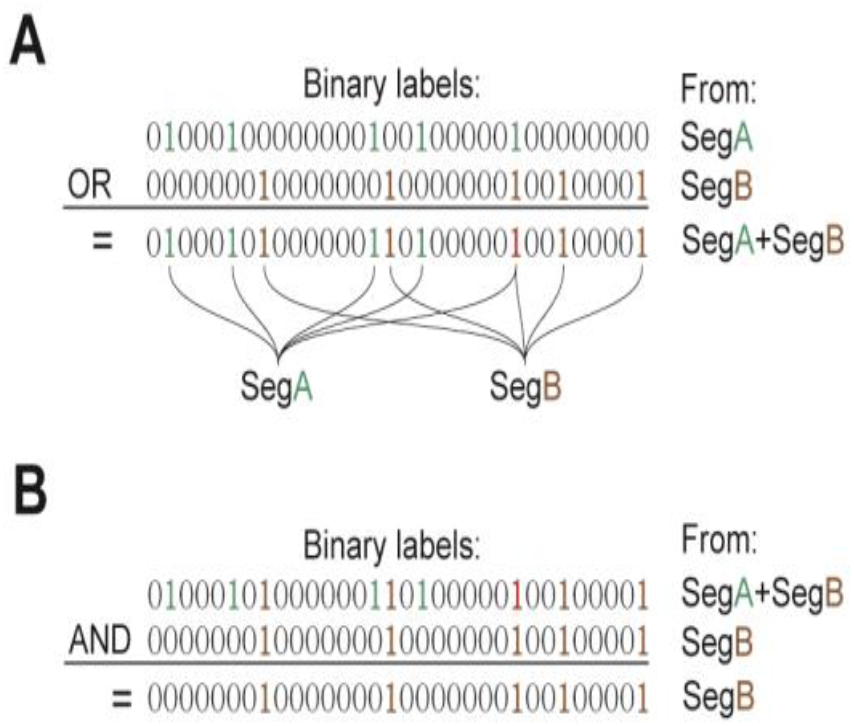
An illustration of the bitwise OR and AND operation used in Aperture. (A) For *k*-mers derived from distinct multiple segments in genome, we merge their binary labels by a bitwise OR operation to keep the segmental positions given by all input labels. The bit in the resulting binary representation is 1 if any bit in the compared position is 1; otherwise, the result is 0. (B) A fast approximation of set intersection, implemented by a bitwise AND operation of binary labels, is performed to detect breakpoint of structural variations. If both bits in the compared position are 1, the bit in the resulting binary representation is 1; otherwise, the result is 0.

### *K*-mer libraries building

The first step in Aperture is to build *k*-mer libraries from the reference genome. A *k*-mer is a nucleotide sequence of a certain length, and the default *k* in Aperture is 23. In order to cover all possible orientations of breakpoints, the reverse complementary sequence of each *k*-mer is also stored in the library. In practice, only the first one in every three consecutive *k*-mers is stored in these libraries, achieving efficient use of memory.

In addition to the *k*-mer sequence itself, the genomic position of each *k*-mer is stored in libraries. In Aperture, we divide reference genome into segments ranging from 30,000 bp to 65,000 bp in length, and inspired by Bloom Filter, we randomly assign each segment with a 32-bit binary label containing only five 1-bits. We also record the position in segment as offset for each *k*-mer, which is necessary for restoring exact genomic position. To accommodate genetic variation in *k*-mer based searching, in Aperture, we add *k*-mers covering common single nucleotide polymorphisms (SNPs) in dbSNP database into the 23-mer library.

Next, we sort all *k*-mers in the library using parallel quicksort algorithm and implement deduplication for *k*-mers with multiple positions in genome. For *k*-mers derived from distinct segments in genome, we merge their binary labels by a bitwise “OR” operation to keep the segmental positions given by all input labels. The bitwise “OR” operation sets 1 in each bit position for which the corresponding bits of either or both labels are 1 s, implementing a fast approximation of set union (Fig.2A). In the meanwhile, the offset positions of these repetitive *k*-mers are set to −1 representing invalidation. Then, to evaluate the mapping qualities of breakends, we also score the repeatability of each *k*-mer. 5-point is given for uniquely mapped *k*-mers, 1-point indicating high repeatability is given for *k*-mers with FFFFFFFF hex labels (containing 32 1-bits), and 3-point is given for the rest. In addition, we consider the potential repeatability of unique *k*-mers. 3-point is given for uniquely mapped *k*-mers that collapse with existing repetitive *k*-mers in library by altering any nucleotide.

To enhance precision of mapping of reads involving repeats, we construct two more libraries in Aperture, a 41-mer library and a spaced-seed library for genomic regions covering repetitive 23-mers. 41-mers are capable of dealing with short repeats, but for much longer repetitive sequences, spaced-seeds are utilized to achieve more accurate searching. Spaced-seeds are a modification to the standard *k*-mers, in which some wildcard positions are added. They span larger range of sequence but only take up less memory. Aperture utilizes them to rescue reads containing consecutive repetitive 41-mers and attempt to obtain the exact match in the whole genome. To save memory use, longer *k*-mers are stored in libraries only if they cover repetitive 23-mers.

### Reads filtering, reads merging and *k*-mer based searching

Aperture takes paired-end reads in FastQ format as inputs. First, we filter out low quality reads. Reads with more than 40% bases of low quality (Phred < 20) or containing bases N are discarded. Then, we extract barcoded sequences from ctDNA reads according to input parameters. In order to identify identical ctDNA molecule when the barcode information is absent, we also extract part of the sequence from fragments as artificial barcodes. At last, barcodes from both sides are concatenated together as a combined UMI which is used to identify PCR duplicates in deep sequencing.

Because the fragment size of ctDNAs are shorter than the read length of NGS paried-end sequencing, overlap of the two read in paired reads are common. To trim redundant sequence, 23-mers shared in both reads of a read pair are found using a super-fast strategy called direct addressing, and then paired-ended reads are assembled as a single consensus sequence.

After PE merging, sequences of ctDNA fragments and their reverse complementary forms are decomposed into 23-mers, followed by binary search against 23-mer library. In practice, we optimize the runtime by performing searching for the first *k*-mer in every two consecutive *k*-mers. The binary labels of others can be obtained by adjacent *k*-mers. After 23-mer searching, the corresponding binary label will be attached to uniquely matched 23-mers, along with a repeat score of 5. If continuous poorly mapped 23-mers are found in a fragment, 41-mers and spaced-seeds sequences will be generated from the fragment and searched against the corresponding libraries to perform remapping. This step rescues those 23-mers covering repeated sequences and attempts to align them uniquely to genome. To mark those 23-mers that are remapped by longer *k*-mer libraries, their repeat scores are set to 3 points.

To further optimize the runtime for Aperture, we modified the algorithm in *k*-mer searching. Rather than performing binary search against all *k*-mers in the library, leading to O(logn) time complexity, we utilized hash table to reduce the search range. The first 15 characters in a 23-mer was converted into an integer number as the hash code. Each hash code corresponded to a hash table, and thus 4^15^ ≈ 1,000,000,000 tables were built, which was nearly the total amount of 23-mers in the library. In such design, only one 23-mer was stored in each table on average, leading to nearly a constant time complexity in library searching.

### SV Breakpoint detection

Aperture uses a fast approximation of set intersection approach to perform accurate breakpoint identification. After *k*-mer based searching, each *k*-mer from a NGS read is attached with a binary label indicating the segment it belongs to. Then, we scan all binary labels for a read and check whether they come from different segments, since *k*-mers from distinct genomic position is a strong evidence of structural variations. To achieve this, we iteratively perform the bitwise “AND” operations on the binary labels of *k*-mers in a queue (Fig.2B). After each operation, we examine whether the resulting label is still valid (i.e. contain at least five 1 bits). A breakpoint candidate is marked for a read if the resulting label is no longer valid.

After breakpoint detection, we filter out breakpoints derived from single nucleotide variants (SNV) and small insertions and deletions (InDel) according to the offsets provided by *k*-mers flanking the breakpoints. Then, we calculate the average of repeat scores given by supporting *k*-mers for both sides as the mapping quality for the read. This score ranges from 1 to 5. Additionally, we count the number of *k*-mers that support each end of the breakpoint. Any breakpoint with less than 3 supporting *k*-mers is removed to reduce false positives.

### Candidates clustering

After gathering all candidates, breakpoints sharing identical junction sequences or similar positions need to be clustered together. First, we sort all candidates by segments of the 5’ end. Then, within each segment, we cluster and merge candidates sharing identical or part of junction sequence constructed from *k*-mers covering the breakpoints. However, small variants and errors near breakpoint may alter junction sequence, leading to failure of clustering. In order to tolerate sequencing errors, PCR errors and mutations, we cluster candidates with similar genomic positions. First, candidates within each segment are sorted according to their offsets of 5’ ends. Then, an overlapping rate of two junction sequences are calculated to quantify the similarity of different candidates. If two candidates are quite similar (overlapping rate <0.2) and very close to each other (within 100 bp), they are clustered together. In addition, the similarity of 3’ end of the breakpoint and reference is checked to avoid false positives result from variants near the breakpoint junction (marked as FAKE_BP in the filter). To calculate the overlapping rate, we first break two junctions into 11-mers and acquire overlapping 11-mers to obtain their subsequences. Then, the overlapping rate can be calculated as the ratio of length of the subsequence and size of the shorter junction.

### Filtering for molecular barcoding-based cfDNA sequencing

Before Aperture reports the final list of SVs, several filters were applied to remove false positives. Firstly, we discard SV clusters with limited ctDNA molecule supports and low mapping qualities. To achieve this, we count the number of distinct barcodes as C and count barcodes that appear only once as U. The reliability of a cluster is quantified using the formula R=C-U/2. A cluster can pass this filter only if it meets one of the following criteria: (1) Mapping quality > 7.2, *k*-mer counts for both sides of the breakpoint >= 4, and R >= 2.5 for dual barcodes dataset or R >= 2 for single barcode dataset (2) Mapping quality > 6.7 and R > 5 (3) Mapping quality > 6.2 and R > 7 (4) *k*-mer counts for both sides >= 8, mapping quality > 9.5, C>=2 and split support >= 8.

We define structural variations as genomic alterations that involve translocations and insertion/deletions that are larger than 50 bp. Therefore, we filter out breakpoints that cover two segments that are separated less than 50 bp in genome coordinates. We also remove candidates with no split read supports and candidates marked as FAKE_BP, which are false positives caused by wrong mapping in highly repetitive regions.

### Simulation of ctDNA datasets

To systematically assess the performance of Aperture, we generated a simulated cfDNA dataset with structural variant breakpoints derived from deletions, inversion and translocations. The first step is to generate an *in silico* tumor genome containing structural variants. Since what we simulate is captured data, to ensure that these variants can be targeted, at least the sequence on one side of the breakpoint (breakend) must fall in an exon of a panel gene. Firstly, we selected 509 tumor-associated genes, and then randomly chose 70 and 40 exons as one breakend of deletions and inversions respectively. The exact position of this breakend in the exon was determined randomly, and the other breakend was randomly placed in 2kbp downstream. For translocations, two groups of 140 exons were picked at random as the starting point of two fragments, and lengths of two detached fragments that are both smaller than 1000 bp were chosen randomly. Then RSVSim[15] was utilized to simulate these structural variations in human reference Hg19 as *in silico* tumor, while Venter genome was selected as the normal genome.

The second step is to generate reads from the given genome. Wessim2 [16] was used to simulate targeted sequencing reads after 24751 probes involving 509 cancer-associated genes were designed by Target Capture Probe Design Tool (https://sg.idtdna.com/site/order/ngs). Then, we filtered out breakpoints that neither ends overlap at least 20bp with target regions, and eventually we got 530 breakpoints as ground truth in the following test (Supplementary Table S1). The error model for 150 bp paired-end sequencing was generated based on a set of real ctDNA data using GemErr module in GemSIM v1.6 [17]. The parameter related to fragment size in Wessim2 was set to -f 170 -d 50 -m 150, representing the fragment size, standard deviation and minimum fragment length, respectively. In order to evaluate performance with different ctDNA fractions, we selected 7 dilutions, 10%, 1%, 0.8%, 0.6%, 0.4%, 0.2%, and 0.1%. About 20,400,000 reads (1350x) from Venter genome and about 2300000 reads (150x) from the *in silico* tumor genome are mixed together to achieve simulated test sample in 10% dilution. For other dilutions, 230,000 reads (15x), 181,000 reads (12x), 136,000 reads (9x), 90,000 reads (6x), 45,000 reads (3x) and 23,000 reads (1.5x) from tumor genome were mixed with 23,000,000 reads (1500x) from Venter genome to achieve dilutions of 1%, 0.8%, 0.6%, 0.4%, 0.2% and 0.1%, respectively.

Since Aperture is developed for ctDNA datasets with multiple duplicates, simulation of ultra-deep sequencing need be done before performance test. The distribution of amplification ratios comes from a real ctDNA data (sample A1 in the results), and each simulated read will replicate certain times according to this distribution. In addition, we take first 6 bp of each read as barcode. For other SV callers, simulated reads without replicates were mapping to human reference Hg19 using BWA mem algorithm[18], then we take the sorted bam file as input. Aperture, GRIDSS and SvABA report SV breakpoints directly as BND type in variant call file (VCF), while Lumpy, Delly and SViCT output breakpoints (BND) as well as INV and DEL events. Therefore, it is necessary to convert SV predicts marked as INV and DEL into BND format. Unlike DEL, an INV event must be transformed to two BND records for both strands in VCF file. In addition, Aperture filters out small events and outputs large structural variations (>50bp) in the final results.

### Sequencing of real patient data

We performed duplex sequencing for cfDNA samples derived from real patients, tagging duplex DNA with adapters containing random but complementary double-stranded barcodes prior to PCR amplification[19]. In this study, 10-20 ng of plasma cfDNA was used directly to prepare libraries without fragmentation. After end repair and A-tailing, the DNA fragments were ligated to our custom adapters. To label each ctDNA molecule, we used a mixture of 8 bp index adapters as exogenous barcodes. Then, Adapter-ligated DNA was amplified for 9 cycles with the KAPA HiFi PCR kit. To capture cfDNA molecules of interest, we designed a set of RNA probes involving 63 cancer relative genes and performed the capture experiments using SureSelect XT Kits (Agilent Technologies, USA) following the manufacturer’s instructions. The prepared samples were then sequenced using Illumina HiSeq X producing 151bp paired-end reads (Illumina, USA).

### Preprocessing of real patient data

Aperture accepts dual barcodes and supports ultra-deep sequencing, however, for competing callers, barcodes and multiple duplicates needed to be removed before testing. First, cfDNA reads without barcodes were aligned to human reference Hg38 using BWA mem algorithm [18]. Then, alignments were sorted and indexed by SAMTools [20]. Finally, we performed duplication using REMOVE_DUPLICATES option in MarkDuplicates command in GATK4 and generated BAM files as input for the test.

## Results

### Performance comparison in simulated cfDNA datasets

Using the simulated cfDNA datasets, we systematically compared the performance of Aperture with a cfDNA-tailored SV caller SViCT, as well as five general purpose SV callers including GRIDSS, Lumpy, SvABA, DELLY and CREST. There are 530 breakpoints as true positives in the 10% dilution dataset. We assessed performance of each SV caller in terms of sensitivity and positive predict value (PPV). Sensitivity is calculated as the ratio of the number of true positives and the total number of true positives. PPV is the ratio of the number of true positives and the total predicts. Also, we calculated the F1-score for each caller, which is a combined measure of sensitivity and specificity (calculated as the harmonic mean of PPV and sensitivity).

In the 10% dilution dataset we found that Aperture had both the highest sensitivity (77.5%) and the highest PPV (91.1%) (Fig.3A). Aperture also had the highest F1-score (0.838) (Fig.3B). On the 1% dilution dataset, Aperture also reached the greatest sensitivity and specificity among 7 tools, predicting 249 true positives and achieving the highest F1-score (0.617). Lumpy ranked the second in sensitivity, identified only 81.9% true positives of which Aperture found. (Fig.3A)

**Figure 3.**
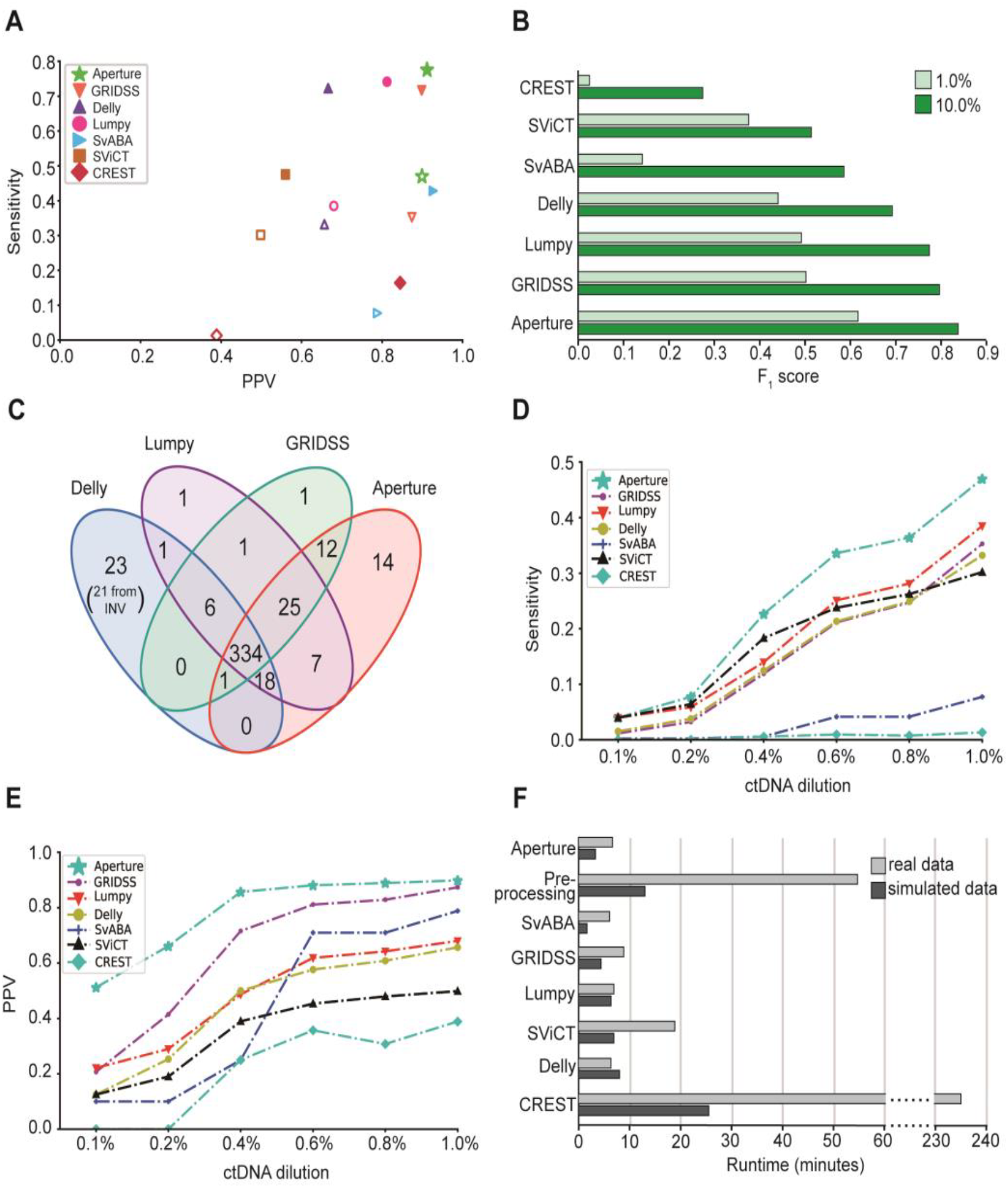
Performance comparison of Aperture with GRIDSS, Delly, Lumpy, SvABA, SViCT and CREST in simulated cfDNA datasets. (A)Comparison of Aperture with other SV callers in terms of sensitivity and positive predictive value (PPV) for the two simulated datasets with different ctDNA dilutions: 10% (solid) and 1% (hollow). (B) Comparison of detection accuracy in 10% and 1% dilutions in term of F1-score, a combined measure of sensitivity and specificity. (C) Structural variation calls reported by four methods in 10% dilution test are presented in a Venn diagram. (D) Comparison of sensitivity as a function of ctDNA dilution for different SV detection methods. (E) Comparison of specificity as a function of ctDNA dilution for different SV detection methods. (F) Total runtime for several SV detection tools in 10% dilution test and a real data test (A1).

In the 10% dilution test, there are 14 true positive breakpoints reported by Aperture only, and half of them are supported by at least 5 ctDNA molecules. We observed that all these high quality predicts are supported by reads containing incomplete breakpoint junctions. Aperture clusters candidates sharing even part of junctions, while other methods ignore these evidences, which may prevent candidates from passing quality filters. This design greatly enhances the sensitivity by rescuing those SVs of which only one breakend falls into the target panel. In addition, we noticed that among the 7 high quality SVs identified by Aperture only, two breakpoints cover repetitive regions and one breakpoint is supported by reads of low mapping qualities, suggesting that repeat might be another barrier for SV detection under the existing approaches. However, a unique strategy of *k*-mer based searching involving three different libraries was utilized in Aperture, allowing to perform more accurate mapping for reads covering repeats, and thus further enhanced performance. In addition, we noticed that Delly managed to identify 23 breakpoints that were missed by other callers. We examined them carefully and found out that 21 records were converted from INV events and were poorly supported. For Delly, identifying these INV events required less supporting reads than predicting both breakpoints (BND) separately in Aperture and GRIDSS (Fig.3C).

To further investigate the performance of Aperture for ctDNAs at very low fraction, we performed tests on 0.8%, 0.6%, 0.4%, 0.2%, and 0.1% dilutions datasets, respectively (detailed results are shown in Supplementary Table S2). Although for all SV callers the sensitivity decreased as ctDNA dilution declined, Aperture remained as the top one in all tests (sharing top sensitivity with Lumpy and SViCT in 0.1% test) (Fig. 2C). In 0.8% and 0.6% dilutions, Aperture was still able to detect more than one-third true positives (36.4%, 33.6%), whereas the sensitivity of GRIDSS, Lumpy, DELLY and SViCT was at least 8 percentage lower. And the sensitivity of SvABA and CREST even fell below 5%. In extremely low dilution (0.2%), Aperture identified 41 true positives, achieving 20% higher sensitivity and 3.5 times higher specificity compared to SViCT, which ranked second in sensitivity in this test (Fig.3D).

In terms of PPV, Aperture also outperformed other SV callers at all seven dilutions (Fig.3E). SV detection became more difficult as the ctDNA fraction dropped below 0.5%. When the ctDNA dilution was above 0.6%, GRIDSS, Lumpy, DELLY and SvABA maintained relatively high PPV (>0.5). However, when ctDNA fraction decreased to 0.2%, all these callers showed a significant decrease in specificity. When the dilution reached 0.1%, only Aperture achieved a PPV of more than 50%. Together, these results suggested that Aperture has an advantage in detecting SVs for cfDNA samples with very low tumor load.

Aperture achieved these results without requiring many computational resources. For the 10% dilution test, Aperture took 204s to complete SV calling under 16 threads, while for real patient sample A1, Aperture consumed 397s to accomplish detection. The run-time for Aperture is shorter than most SV-calling algorithms, even not counting the additional time for data pre-processing that includes alignment and sorting for the other methods (Fig.3F). Tests were performed on a machine with dual Intel Xeon CPU E5-2680 v4.

### Detection performance in reference data

To assess Aperture in real targeted cfDNA sequencing data, we obtained a reference dataset containing HD786 from SRA database [9]. The HD786 sample was a “simulated” cfDNA sample mixed from multiple cell line samples. It was sequenced after capturing by a custom gene panel, and repeated once to create a replicate. There were four verified structural variants in the dataset, but two of them were smaller than 50bp in size that are out of range for Aperture detection. Reads in this dataset did not contain barcode adapters, so we regarded the first 6bp in R1 as barcode in this test. After running the test, we found that Aperture was able to detect breakpoints of *SLC34A2*/*ROS1* and *CCDC6*/*RET* in both replicates.

### Sensitive and specific detection of *ALK* fusion in lung cancer patient for treatment monitoring

To further evaluate the performance of Aperture in real patient data, we obtained a set of real data from a non-small cell lung cancer patient with EML4-ALK fusion. This fusion gene, caused by an inversion of chromosome 2, produces constitutively active *ALK* protein, which in turn promote and maintain the malignant behavior of the cancer cells. Patients with the EML4-ALK fusion are treated with *ALK* inhibitors, such as Crizotinib [21] to inhibit tumor growth. In this study, we monitored a patient from receiving Crizotinib as treatment to developing resistance, and collected five cfDNA samples at different time points in the process. All samples were performed deep sequencing after adding dual-index barcoded adapters for each molecule. Aperture accepts raw data as input, however, for other SV callers, alignment file after removing barcodes and multiple duplicates is accepted. Hg38 was used as reference genome in this test.

Sample A1 was collected shortly after being diagnosed with EML4-ALK positive lung cancer, confirmed by NGS of pleural effusion from the patient. At the same time the patient began to receive Crizotinib treatment. Aperture, Lumpy, Delly and SViCT managed to identify the fusion breakpoint in this sample, while GRIDSS missed it since it did not pass their filters (Table 1). SvABA and CREST failed to report this variant even in unfiltered results. In sample A2, which was taken 20 days after crizotinib treatment, Aperture, Lumpy and SViCT were still able to detect this fusion. Results given by Aperture indicated that fraction of this fusion in cfDNA was significantly reduced, most likely due to the treatment of ALK inhibitor. Compared to Lumpy and SViCT, the number of total predicts given by Aperture is about 85% and 82% less, indicating a much higher specificity in detecting low frequency ctDNA SVs. All the algorithms failed to detect this fusion in sample A3, suggesting that under treatment, the amount of tumor cells carrying ALK fusion is very rare in plasma. In sample A4, Aperture identified a novel EML4-ALK fusion (*ALK* exon 20 fused with *EML4* exon 1), but all competing callers failed to detect this variant. Sample A5 was collected after drug resistance occurred, and as expected no algorithm could detect EML4-ALK fusion in this sample. For specificity, we noticed that Aperture achieved the least total predictions among all the callers that managed to identify the *ALK* fusion (Aperture, Lumpy, Delly and SViCT).

**Table 1.**
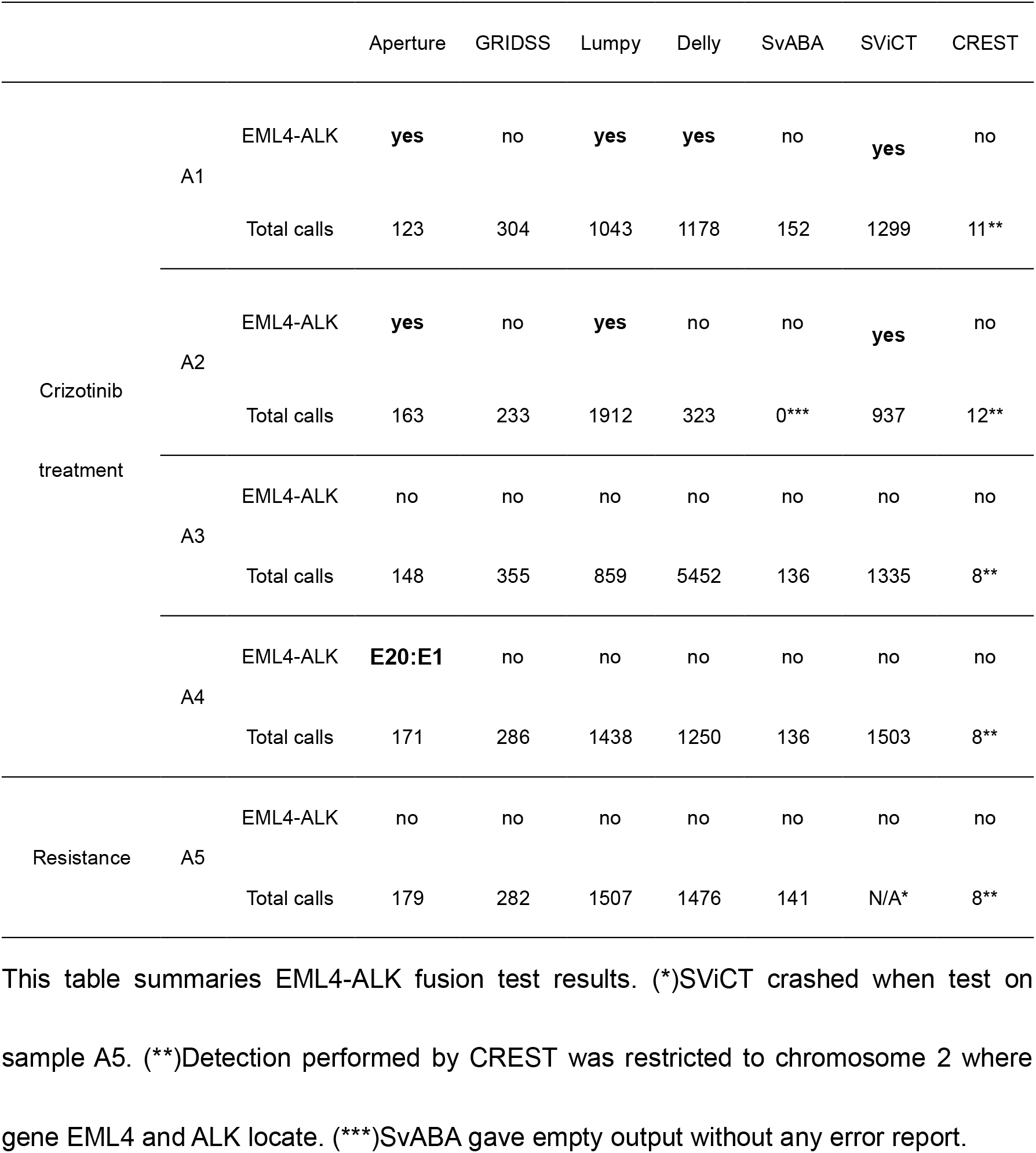
Detection of EML4-ALK translocation by Aperture and compared tools in samples from a lung cancer patient

### Detection of complex *NTRK* fusion in lung cancer patient samples

*NTRK* fusion gene has recently emerged as targets for cancer therapy [22], detecting *NTRK* fusion in ctDNA may help treatment selection when surgery is not available. To assess the ability for Aperture to detect SVs involving *NTRK* genes, we used cfDNA datasets of plasma sample from two lung cancer patients (Table 2). In sample B1, Aperture, GRIDSS, Lumpy, DELLY, SvABA and SViCT all managed to identify the TPR-NTRK1 fusion. In sample B2, only Aperture and GRIDSS were able to identify LMNA-NTRK1 fusion, and Aperture gave the least total predictions. LMNA-NTRK1 had five supporting molecules in the dataset, which should be enough to pass filter in most SV callers. By extracting reads involved by this fusion and performing global alignment by blastN (https://blast.ncbi.nlm.nih.gov/Blast.cgi), we found that the reads covering the NTRK1-LMNA translocation contained a 9bp novel insertion and a 30bp repeated sequence (Fig.4). It is possible that SV callers missed the read due to the low alignment quality given by the aligners for repetitive sequences. In addition, *LMNA* is not included in the targeted panel, so the coverage of the supporting reads are much lower, further increasing the difficulty of identifying this variant. These results suggest that the alignment-free strategy Aperture used can augment the sensitivity for detecting complex SVs involving repetitive sequences.

**Figure 4.**
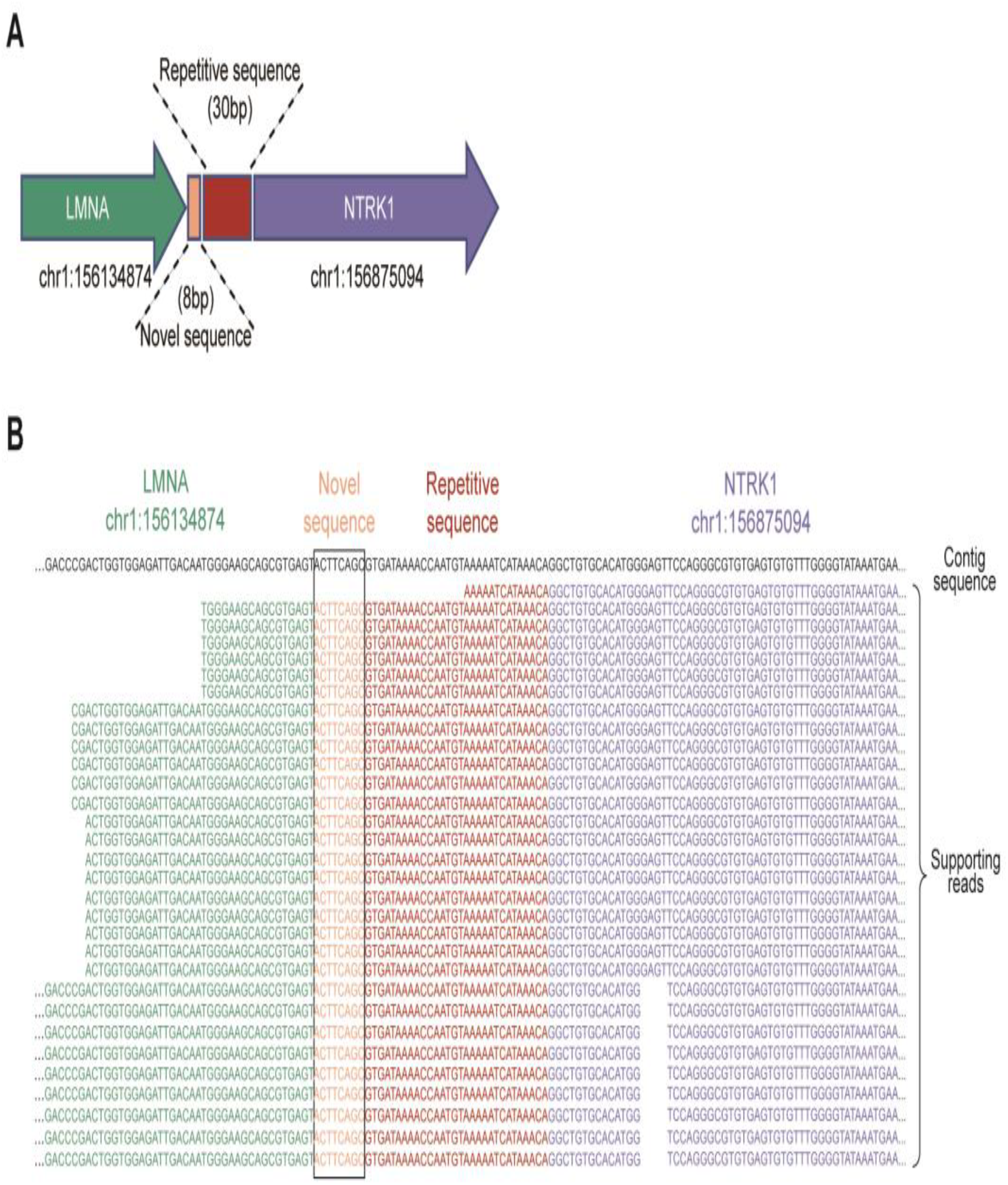
Aperture identifies a complex breakpoint with insertion of a short repetitive sequence. (A) Illustration of the complex SV in chromosome 1 that leads to the fusion of LMNA and NTRK1 gene regions. The breakpoint junction contains a 30-bp repetitive sequence and an 8-bp novel sequence. (B) View of the 31 cfDNA fragments from 5 molecules supporting the existence of the breakpoint junction illustrated in (A).

**Table 2.** Detection of *NTRK* fusion by Aperture and compared tools in two lung cancer patient samples.

|  |  | Aperture | GRIDSS | Lumpy | Delly | SvABA | SViCT |
| --- | --- | --- | --- | --- | --- | --- | --- |
| B1 | TPR-NTRK1 | yes | yes | yes | yes | yes | yes |
|  | Total calls | 58 | 200 | 1526 | 134 | 41 | 952 |
| B2 | LMNA-NTRK1 | yes | yes | no | no | no | no |
|  | Total calls | 112 | 263 | 2490 | 166 | 112 | 2558 |

This table summaries *NTRK1* fusion test results.

### Detection of HBV integration near *TERT* gene in liver cancer patient cfDNA samples

Recurrent HBV integration near cancer-related gene might play an important role in tumorigenesis [12]. To assess the performance of Aperture in detection of viral integration, a special category of structural variation, we used plasma samples from three patients with hepatocellular carcinoma that having HBV integrations near the *TERT* gene. The cfDNA datasets were generated with a targeted panel containing *TERT* promoter. Unlike genomic SV detection, viral genome needs to be included in reference sequences before detection of viral integration.

According to the test results, Aperture, Lumpy and Delly were capable of identifying HBV integrations near the *TERT* gene (Table 3). Lumpy and Aperture both identified the HBV-TERT predictions in sample C1 and C2, but Lumpy gave five more predictions that possess imprecise breakpoints in sample C3. DELLY missed one HBV-TERT predict in sample C1 and C2 and was unable to identify precise HBV-TERT call in sample C3. SvABA and GRIDSS fail to identify any of the HBV-TERT viral integrations. All these integrations failed to pass filters in GRIDSS and marked as LOW_QUA or NO_ASSEMBLY. SViCT does not support viral detection because we noticed small genome such as HBV was missing in VCF header, suggesting that small references might be discarded in SViCT before detection.

**Table 3.**
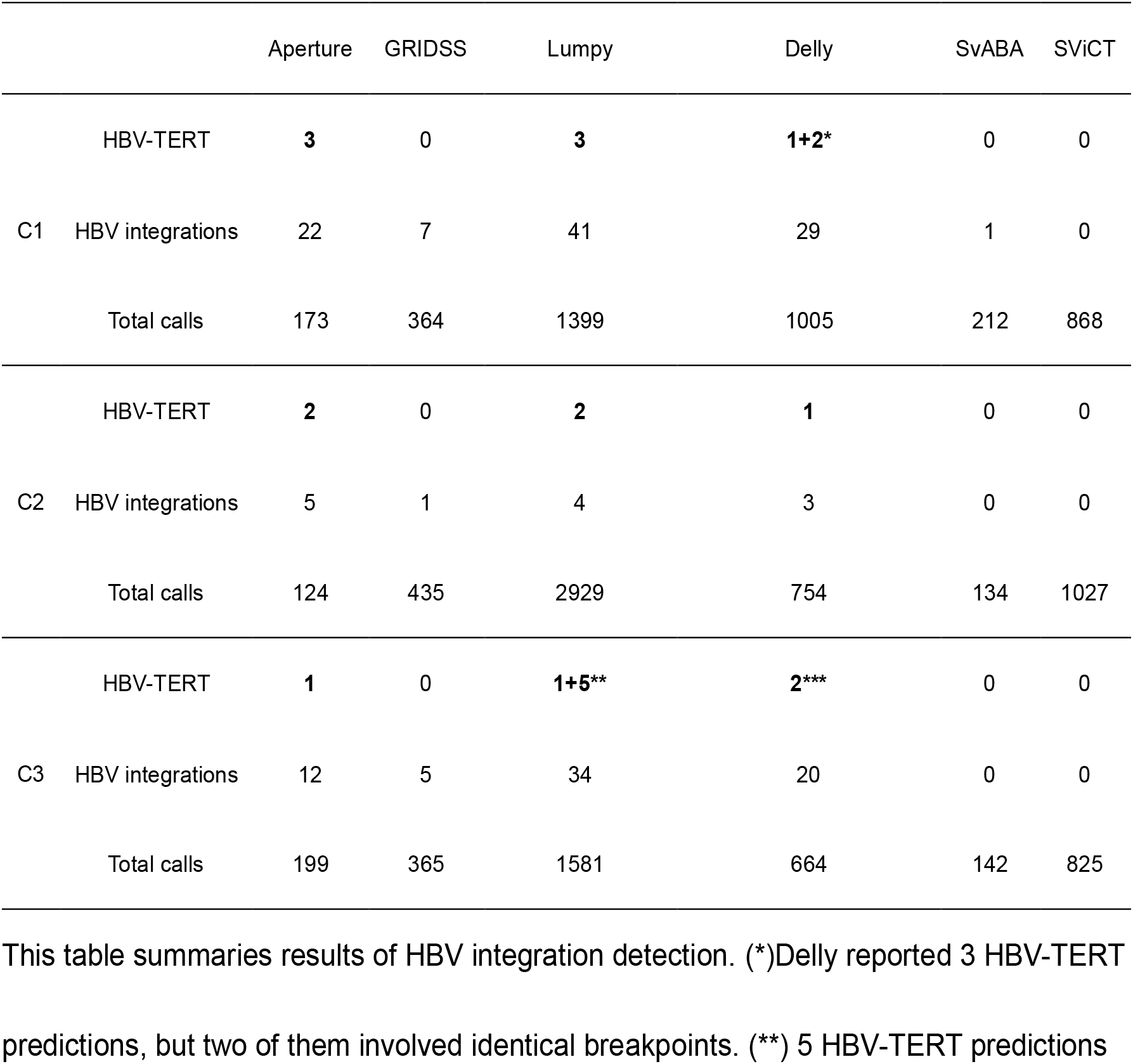
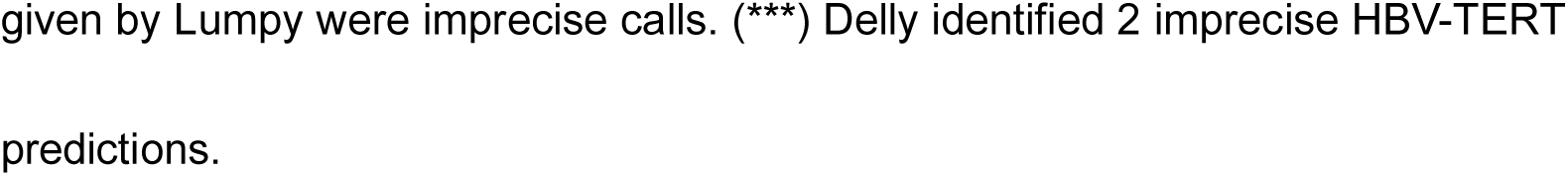
Detection of integration of HBV genome near *TERT* gene by Aperture and five other tools in three liver cancer patient samples.

## Discussion

In this study, we developed a new structural variation and viral integration detection tool for ctDNA dataset. Aperture uses *k*-mer based searching and a fast approximation of intersection to detect potential breakpoints. In addition, Aperture performs candidates clustering to collect evidences efficiently and filters low quality candidates based on counts of ctDNA molecule. With these features, Aperture outperforms existing SV callers in sensitivity and specificity in ctDNA dataset, and is capable of detecting complex SV junctions with low coverage and repeat insertion. Moreover, using parallel programming and *k*-mer based searching, Aperture achieves great efficiency.

CtDNAs are highly fragmented DNAs in bloodstream, with small fragment size between 50 to 166bp [1], making the precise alignments for both sides of ctDNA reads containing SV breakpoints more difficult. Alignment-based callers may yield many false positives owing to sequencing errors or read mis-alignments, especially within repetitive sequences [23]. To enhance accuracy in SV detection in ctDNA, in Aperture, we implemented a unique *k*-mer searching strategy for breakpoint detection. Unlike the single *k*-mer library used in ChimeRScope [24], we built three *k*-mer libraries with different *k*-mer sizes for searching. If a read covers repetitive regions and shorter *k*-mers that fail to map, our algorithm will remap it with longer *k*-mers libraries including 41-mers and spaced-seeds to attempt accurate alignment. Given that aligner may be disturbed by mismatch especially in repetitive regions, we also incorporated 23-mers embedding common SNPs into library to further improve precision for reads containing known variants.

We aimed to implement the detection of breakpoints involving repetitive sequences by *k*-mer based strategy, which is much complicated than fusion gene discovery in RNA sequencing data. Instead of assigning a list of all possible sources for *k*-mers derived from distinct regions, which is memory consuming especially for repetitive regions, we developed a unique fixed size binary label system with a fast approximation of intersection operations. With the system, Aperture is able to detect breakpoints that overlap with novo-*k* mers and repetitive *k*-mers, which can be seen as an extension of the novoBreak method [25]. NovoBreak only detects breakpoints covering novo-kmers. However, in some circumstances, novo-kmers may be absent in breakpoint junction. Typically, there should be at least k novo-kmers in a junction, but for breakpoints covering repetitive sequence, their counts will dramatically decrease. When the repetitive sequence is longer than *k*, no novo-kmers but repetitive *k*-mers can be identified in a junction. In our approach, repetitive sequences that mapped to multiple genomic locations will be given a recalculated binary label according to the binary labels of all the locations it mapped to using a bitwise “OR” operation. For a breakpoint junction involving repetitive sequence, Aperture can jump across this repetitive junction, and stop until a new location is identified by checking the validation of binary labels after each bitwise “AND” operation. Indeed, the tests on both simulated data and real data validated the superior performance of Aperture in comparison to existing methods, especially on the complex SVs containing repetitive regions. Additionally, the number of 1-bits in a valid label is determined by the size of segments divided from genome. To achieve one-to-one mapping of binary labels and segments, we need to generate at most 100,000 valid labels, as the ratio of human genome size (3 billion bp) and minimum length of segment (3,000 bp). Selecting five 1-bits in a 32-bit label can cover 201,376 possibilities, which is just adequate to symbolize all segments in human genome. More 1-bits may lead to higher saturation.

Assembly-based algorithm is widely used in SV callers, because longer contig assembled from short reads can be more accurately mapped to the genome, enabling more sensitive detection of SVs. However, given that ctDNA is often very short, about 166bp, the assembly-based strategy may not be suitable for ctDNA dataset. Too many short and similar sequences and a lack of PE evidences may bring obstacles to obtain longer and correct contigs, resulting in failure to call true variants especially for SVs involving repeated sequence. In addition, sequence assembly has large computational requirements. Therefore, in Aperture, we detect breakpoint independently on each read and perform candidates clustering to merge candidates sharing identical or part of junction sequences. This strategy achieves better sensitivity as well as higher efficiency than other assembly based callers according to the performance tests on both simulated data and real data.

The advantage of Aperture is also evident on detection of viral integration, which plays a critical role in cancer development. Due to the widespread HBV integrations in the genome, we found existing SV callers based on the genome-wide assembly strategy may fail to correctly assemble the breakpoint junction, leading to the removal of true integrations.

How could Aperture’s performance be further improved? First, native support for read sequencing quality could be incorporated into SV filter, which is expected to reduce more false positives and further promote specificity. Second, SV discovery in cfDNA data is limited by the read length. Aperture achieves accurate detection of deletion, inversion and translocation variants, however, for large inversions or complex rearrangements with exceedingly long junction sequence, Aperture may be less sensitive due to the lack of uniquely mapped *k*-mers. Here, we can avoid missing some large SVs by generating longer sequence after merging pair-ended reads. The improvement with the paired-end merging may be not very significant for cfDNA datasets due to the limited length of cfDNA molecules in bloodstream.

## Conclusions

The software Aperture presented here achieves highly sensitive and specific detection of SVs and viral integrations for ctDNA dataset, even when the SVs involve complex junctions. Our work may enhance the diagnostic potential of ctDNA in early cancer detection and help in treatment monitoring, for some fusions and viral integrations are closely related to certain tumor types. In addition, Aperture runs fast and is efficient in consumptions of computational resources. For these reasons, we believe that the method proposed in this study will not only be helpful in bioinformatics community, but also offer a reliable tool in research and clinical practice and help progress precision medicine.

## Availability and requirements

Project name: Aperture

Project home page: https://github.com/liuhc8/Aperture

Operating system(s): Platform independent

Programming language: Java

Other requirements: Java 1.8.0 or higher

License: Apache-2.0 License

## Abbreviations

ctDNA: circulating tumor DNA
cfDNA: cell-free DNA
SV: structural variation
NGS: next-generation sequencing
SNV: single nucleotide variants
InDel: insertions and deletions
VCF: variant call file
PPV: positive predict value
PE: paired-end
SNP: single nucleotide polymorphism

## Declarations

### Ethics approval and consent to participate

Plasma samples from patients were obtained under the protocols approved by the Institutional Review Board at Cancer Hospital of Chinese Academy of medical sciences, with informed consent for research use.

## Consent for publication

Not applicable.

## Availability of data and materials

Reference datasets, from a previous publication, were downloaded from NCBI Sequence Read Archive (SRA) under accession number SRR8551544 and SRR8551545.

The executable file, source code and the simulation datasets are available at https://github.com/liuhc8/Aperture.

## Authors’ contributions

HL and XW developed the algorithm. HL implemented and tested the code, created the simulated dataset, and ran the comparison tests. GL assisted in algorithm design and ran competing algorithms. JL planned and performed the clinical part of the study. HY collected the clinical samples. HL and XW wrote the manuscript. XW conceived, coordinated and supervised the project. All authors read and approved the final manuscript.

## Acknowledgements

The authors would like to thank Qianqian Song, Xiaofei Zhao and Jie Yang for helpful discussions.

## Reference

1. Wan JCM, Massie C, Garcia-Corbacho J, Mouliere F, Brenton JD, Caldas C, Pacey S, Baird R, Rosenfeld N: Liquid biopsies come of age: towards implementation of circulating tumour DNA. Nat Rev Cancer 2017, 17:223-238.

2. Phallen J, Sausen M, Adleff V, Leal A, Hruban C, White J, Anagnostou V, Fiksel J, Cristiano S, Papp E, et al: Direct detection of early-stage cancers using circulating tumor DNA. Sci Transl Med 2017, 9.

3. Newman AM, Lovejoy AF, Klass DM, Kurtz DM, Chabon JJ, Scherer F, Stehr H, Liu CL, Bratman SV, Say C, et al: Integrated digital error suppression for improved detection of circulating tumor DNA. Nat Biotechnol 2016, 34:547–555.

4. Cameron DL, Schroder J, Penington JS, Do H, Molania R, Dobrovic A, Speed TP, Papenfuss AT: GRIDSS: sensitive and specific genomic rearrangement detection using positional de Bruijn graph assembly. Genome Res 2017, 27:2050–2060.

5. Layer RM, Chiang C, Quinlan AR, Hall IM: LUMPY: a probabilistic framework for structural variant discovery. Genome Biol 2014, 15:R84.

6. Wala JA, Bandopadhayay P, Greenwald NF, O’Rourke R, Sharpe T, Stewart C, Schumacher S, Li Y, Weischenfeldt J, Yao X, et al: SvABA: genome-wide detection of structural variants and indels by local assembly. Genome Res 2018, 28:581–591.

7. Rausch T, Zichner T, Schlattl A, Stutz AM, Benes V, Korbel JO: DELLY: structural variant discovery by integrated paired-end and split-read analysis. Bioinformatics 2012, 28:i333–i339.

8. Wang J, Mullighan CG, Easton J, Roberts S, Heatley SL, Ma J, Rusch MC, Chen K, Harris CC, Ding L, et al: CREST maps somatic structural variation in cancer genomes with base-pair resolution. Nat Methods 2011, 8:652–654.

9. Gawronski AR, Lin YY, McConeghy B, LeBihan S, Asghari H, Kockan C, Orabi B, Adra N, Pili R, Collins CC, et al: Structural variation and fusion detection using targeted sequencing data from circulating cell free DNA. Nucleic Acids Res 2019, 47:e38.

10. De Coster W, Van Broeckhoven C: Newest Methods for Detecting Structural Variations. Trends Biotechnol 2019, 37:973–982.

11. Mahmoud M, Gobet N, Cruz-Davalos DI, Mounier N, Dessimoz C, Sedlazeck FJ: Structural variant calling: the long and the short of it. Genome Biol 2019, 20:246.

12. Jhunjhunwala S, Jiang Z, Stawiski EW, Gnad F, Liu J, Mayba O, Du P, Diao J, Johnson S, Wong KF, et al: Diverse modes of genomic alteration in hepatocellular carcinoma. Genome Biol 2014, 15:436.

13. Snyder MW, Kircher M, Hill AJ, Daza RM, Shendure J: Cell-free DNA Comprises an In Vivo Nucleosome Footprint that Informs Its Tissues-Of-Origin. Cell 2016, 164:57–68.

14. Ma B, Tromp J, Li M: PatternHunter: faster and more sensitive homology search. Bioinformatics 2002, 18:440–445.

15. Bartenhagen C, Dugas M: RSVSim: an R/Bioconductor package for the simulation of structural variations. Bioinformatics 2013, 29:1679–1681.

16. Kim S, Jeong K, Bafna V: Wessim: a whole-exome sequencing simulator based on in silico exome capture. Bioinformatics 2013, 29:1076–1077.

17. McElroy KE, Luciani F, Thomas T: GemSIM: general, error-model based simulator of next-generation sequencing data. BMC Genomics 2012, 13:74.

18. Li H, Durbin R: Fast and accurate long-read alignment with Burrows-Wheeler transform. Bioinformatics 2010, 26:589–595.

19. Kennedy SR, Schmitt MW, Fox EJ, Kohrn BF, Salk JJ, Ahn EH, Prindle MJ, Kuong KJ, Shen JC, Risques RA, Loeb LA: Detecting ultralow-frequency mutations by Duplex Sequencing. Nat Protoc 2014, 9:2586–2606.

20. Li H, Handsaker B, Wysoker A, Fennell T, Ruan J, Homer N, Marth G, Abecasis G, Durbin R, Genome Project Data Processing S: The Sequence Alignment/Map format and SAMtools. Bioinformatics 2009, 25:2078–2079.

21. Christensen JG, Zou HY, Arango ME, Li Q, Lee JH, McDonnell SR, Yamazaki S, Alton GR, Mroczkowski B, Los G: Cytoreductive antitumor activity of PF-2341066, a novel inhibitor of anaplastic lymphoma kinase and c-Met, in experimental models of anaplastic large-cell lymphoma. Mol Cancer Ther 2007, 6:3314–3322.

22. Amatu A, Sartore-Bianchi A, Siena S:NTRK gene fusions as novel targets of cancer therapy across multiple tumour types. ESMO Open 2016, 1:e000023.

23. Ding L, Wendl MC, McMichael JF, Raphael BJ: Expanding the computational toolbox for mining cancer genomes. Nat Rev Genet 2014, 15:556–570.

24. Li Y, Heavican TB, Vellichirammal NN, Iqbal J, Guda C: ChimeRScope: a novel alignment-free algorithm for fusion transcript prediction using paired-end RNA-Seq data. Nucleic Acids Res 2017, 45:e120.

25. Chong Z, Ruan J, Gao M, Zhou W, Chen T, Fan X, Ding L, Lee AY, Boutros P, Chen J, Chen K: novoBreak: atlocal assembly for breakpoint detection in cancer genomes. Nat Methods 2017, 14:65–67.

